# Anatomically and Physiologically Accurate Engineered Neurovascular Unit and Blood-Brain Barrier Model Using Microvessels Isolated from Postmortem Human Brain Tissue

**DOI:** 10.1101/2024.09.26.615283

**Authors:** Brian J. O’Grady, A. Scott McCall, Samuel Cullison, Daniel Chavarria, Andrew Kjar, Matthew S. Schrag, Ethan S. Lippmann

## Abstract

Brain vasculature is a complex and heterogeneous physiological structure that serves specialized roles in maintaining brain health and homeostasis. There is substantial interest in developing representative human models of the brain vasculature for drug screening and disease modeling applications. Many contemporary strategies have focused on culturing neurovascular cell types in hydrogels and microdevices, but it remains challenging to achieve anatomically relevant vascular structures that have physiologically similar function to their *in vivo* counterparts. Here, we present a strategy for isolating microvessels from cryopreserved human cortical tissue and culturing these vessels in a biomimetic gelatin-based hydrogel contained in a microfluidic device. We provide histological evidence of arteriole and capillary architectures within hydrogels, as well as anastomosis to the hydrogel edges allowing lumen perfusion. In capillaries, we demonstrate restricted diffusion of a 10 kDa dextran, indicating intact passive blood-brain barrier function. We anticipate this bona fide human brain vasculature-on-a-chip will be useful for various biotechnology applications.

## Introduction

The human brain’s intricate physiology is supported by a complex network of vasculature spanning a heterogeneous arteriovenous axis.^1^ The brain vasculature carries out diverse functions and possess unique cellular and molecular signatures relative to peripheral vasculature. For example, arterioles contribute to blood flow regulation through neurovascular coupling mechanisms,^2^ while capillaries serve as the site of material exchange between the bloodstream and parenchyma, as controlled by the properties of the blood-brain barrier (BBB).^3^ Anatomically, vessels in the brain are lined with highly specialized endothelial cells and ensheathed by mural cells within a collagen-rich basement membrane. The outer edge of the vessels is surrounded by astrocyte endfeet, collectively creating the intricate three-dimensional structure of the neurovascular unit (NVU). All of these cells actively communicate to ensure proper neurovascular function, and in various disease states, these functions are compromised.^4^ As such, there is significant interest in developing more physiologically relevant *in vitro* models of the human cerebrovasculature to better understand disease mechanisms and develop intervention strategies.

Many strategies have been developed to recapitulate the three-dimensional architecture of the NVU *in vitro*.^5–7^ These strategies generally rely on the self-assembly capacity of endothelial, mural, and glial cells, which are typically sourced from primary material or differentiated from human pluripotent stem cells (hPSCs). In general, cells are cultured in hydrogels contained within well plates or microfluidic devices, the latter of which permits assessments of vascular perfusion and measurements of barrier properties. Culture in hydrogels allows growth or dynamic remodeling as neurovascular cells assemble into their desired structures.^8,9^ Other models have relied on direct placement of cells in desired compartments, for example lining prefabricated channels in hydrogels with endothelial and mural cells, while astrocytes or other neural cells are cultured within the hydrogel.^10–12^ While all of these models have achieved some structural features of cerebral vasculature, they generally lack the organized architectures seen *in vivo*, which would ultimately be more desirable for studying the intimate communication between cell types in the NVU across different levels of the vascular tree.

Here, to better mimic the endogenous NVU and cerebrovasculature architectures *in vitro*, we present a model that is fabricated using microvessels enriched from postmortem, cryopreserved human cortical tissue (Figure 1). We developed an enzyme-free method to collect and purify human microvessels, which helps maintain native vessel architectures. Then, rather than relying on assembly of individual cell types into the desired neurovascular structures, these pre-existing vessels are embedded in a peptide-functionalized gelatin-based hydrogel within a custom microfluidic chip under conditions that promote vascular growth. Using immunohistology and live imaging, we demonstrate that these vessels possess native architectures, are perfusable, and retain cellular identities reminiscent of cerebral vasculature. Excitingly, interconnected arterioles and capillaries are reliably detected in the hydrogels, and the capillaries demonstrate robust passive BBB function as indicated by a tracer extravasation assay. Overall, we anticipate this model system will open new avenues for understanding neurovascular function in health and disease.

**Figure 1:**
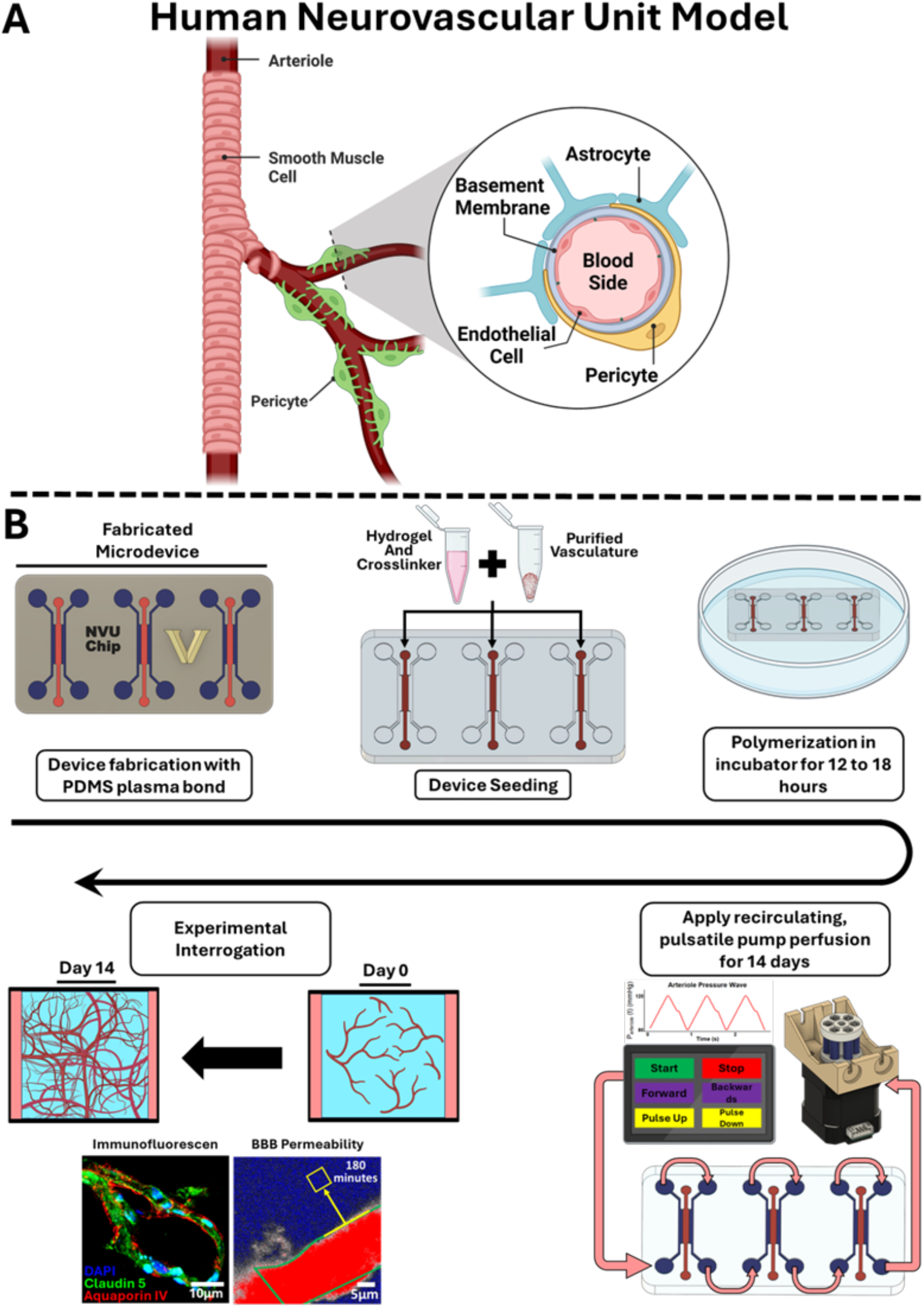
Overview of the *ex vivo* human neurovascular model. **A)** A detailed graphic representation of the human neurovascular unit (NVU), highlighting the spatial arrangement of lumenized endothelial cells surrounded by a basement membrane, pericytes, and astrocyte endfeet. **B)** Timeline of the NVU model fabrication and experimentation process. Custom microdevices are fabricated and seeded with purified cerebral blood vessels in an enzymatically crosslinked hydrogel. The hydrogel undergoes polymerization in an incubator for 12 to 18 hours, and then a recirculating pulsatile pump perfuses the microdevice for 14 days. During this period, the vasculature grows and develops within the hydrogel matrix. On day 14, various analyses are performed to assess the functionality and representativeness of the NVU model.

## Methods

### Microfluidic chip design, fabrication, and 3D printed tools

The master mold for the microfluidic chips were fabricated either using a stereolithographic (SLA) 3D printer (Formlabs 3) or a Haas CM-1 CNC. Printed molds were fabricated using Formlabs Black resin v3. The molds were extensively washed in 100% isopropanol alcohol to remove all uncrosslinked monomers and photocrosslinking agents. The molds were then extensively washed with ultra-pure water and allowed to air dry in front of a fan for 30 minutes. The washed molds were then placed in a Formlabs Cure chamber and exposed to ultraviolet light for 10 minutes. All prints were then treated with parylene as previously described,^13^ which allows them to serve as master molds for elastomers. In some cases, 6061 aluminum stock (9146T69; McMaster-Carr) was milled on the CNC to create master molds. Additional information (including CAD files, tools used, generated paths, feeds and speeds, etc.) are available on GitHub.

To create microdevices, PDMS (Sylgard 182; Dow Corning) was poured onto the 3D printed and/or CNC machined master molds, and the molds were then placed in a vacuum chamber to remove all bubbles and subsequently heat-cured at 80°C for 30 minutes. After heat-curing, the solidified PDMS was carefully removed from the master molds and a biopsy punch (1.5 mm) was used to create inlets and outlets for the hydrogel injection channel and the media perfusion channels. Using scotch tape, debris was removed from the surface of the PDMS, and then microdevices were plasma bonded to a precleaned microscope glass coverslip (Ted Pella 260461-100). Prior to experimentation, microdevices were sterilized using an autoclave. STL files and parts lists for all microdevices are available on GitHub.

### Pump perfusion system

A customized pump perfusion system was modified from a previous system.^14^ Briefly, the system uses a Raspberry Pi 4, Raspberry Pi touchscreen, Easydriver, and a stepper motor with a 12v power supply. A low-cost, custom printed circuit board was designed and fabricated to integrate the electronics for this system. Custom code was written to generate desired pulse waves, where the software allows the user to control the direction of the fluid flow, duration to the systolic amplitude, pulse duration, time in between pulses, duration of the dicrotic notch, and control over the timing of the dicrotic notch. Schematic designs, CAD files, device assembly, design rule checking list, all code, design files, bill of materials, and assembly videos are available on Github.

### GelCad biomaterial synthesis

GelCad biomaterial was synthesized as previously described with minor modifications.^15,16^ Briefly, Type A porcine skin gelatin (Sigma) was reconstituted in triethanolamine (TEOA; Sigma) to create a 10% w/v solution and stirred at 37 °C for 2 hours until fully dissolved. The pH of the solution was then adjusted to 8.0–8.5 by adding 1.0 M HCl or 1.0 M NaOH. An intermediate reaction was used to prepare the peptides for binding. 80 mg of ethylcarbodiimide hydrochloride (EDC; Thermo Fisher) and 120 mg of n-hydroxysuccinimide (NHS; Thermo Fisher) were mixed with 80 mg of peptide. The combined solids were dissolved in 10 mL of N–N-dimethyformamide (DMF) (28 mg/mL), and 15 mL of PBS (19 mg/mL) at 37°C for 1 hour. The peptide intermediate was then added dropwise to the gelatin solution and reacted for 4 hours at 37°C with constant stirring. Next, the reaction was quenched by raising the pH to 8 using sodium bicarbonate (5 M). The final solutions were dialized using a Slide-A-Lyzer Dialysis Flask (ThermoFisher 87761; 3.5 kDA MW cuttoff) with ultrapure water. After filtration, the biomaterial solutions were frozen, lyophilized, and pulverized for long term storage.

### Complete media preparation

Heat stable basic fibroblast growth factor (bFGF; Gibco PHG0367) was reconstituted by adding 1 mL of PBS to a 5 μg vial. Epidermal growth factor (EGF; PeproTech AF-100-15-1MG) was reconstituted by adding 1 mL of PBS to a 1 mg vial. Fibronectin (ThermoFisher 33016015) was reconstituted by adding 5 mL of PBS to a 5 mg vial. Vascular endothelial growth factor (VEGF, PeproTech 100-20) was reconstituted by adding 1 mL of PBS to a 10 μg vial. Thymosin Beta-4 (TB4, PeproTech 140-14) was reconstituted by adding 1 mL of PBS to a 20 μg vial. All components were aliquoted and stored at -20°C. Complete media was prepared by combining 17.75 mL of human endothelali serum-free medium (hESFM; ThermoFisher 1111044) with 1 mL reconstituted bFGF, 10 μL reconstituted EGF, 500 μL reconstituted fibronectin, 500 μL reconstituted VEGF, and 250 μL reconstituted TB4, creating a final volume of 20 mL.

### Microfluidic chip design

The microfluidic chip features a central channel measuring 1.532 mm in width and 0.152 mm in depth, flanked by two larger perfusion channels, each 1.047 mm in width. This configuration accommodates large fluid volumes. The chip’s pillarless design eliminates the traditional structural supports that can lead to hydrogel spillover into perfusion channels—a common issue in larger-featured devices. This design enhances the reproducibility of the experimental setup by preventing asymmetry in the chips, which could otherwise compromise experimental outcomes. Furthermore, the central channel’s extended length offers enhanced control during the hydrogel injection process. This design feature further ensures that the hydrogel remains confined to the designated area, minimizing the risk of leakage into adjacent perfusion channels.

### Microfluidic chip holder for hydrogel injection

A custom-designed microfluidic chip holder, depicted in Supplemental Figure 1, was developed to complement the chip design. This holder positions the chip at a precise angle during hydrogel injection, which helps fill the central channel without spillage of hydrogel into the perfusion channels.

### Collection of human brain tissue and preparation for microvessel enrichment

Human brain tissue was obtained at autopsy from a 73-year-old male patient who presented with mild to moderate generalized atrophy, severe Alzheimer’s disease (Braak stage IV, Thal stage 5, A3, B3, C3), moderate cerebral amyloid angiopathy (CAA), and mild atherosclerosis. There was no evidence of Lewy bodies or other tauopathies. The brain weight was recorded at 1,230 grams, and the post-mortem interval was 8 hours. Tissue collection was conducted under an IRB-approved protocol at Vanderbilt University Medical Center (#180287). After manually separating cortical grey matter from white matter, tissue was cut into ∼1 cm^3^ pieces and placed into cryovials that were subsequently filled with BrainBits Hibernate A media containing 10% DMSO. Cryovials were slowly cooled in a Nalgene Mr. Frosty container overnight in a -80°C freezer. The following day, cryovials were placed in liquid nitrogen for long-term storage. For experiments, a single cryovial was removed from liquid nitrogen and slowly brought up to room temperature in a liquid or bead bath. Once 70% thawed, the vial was transferred to a sterile cell culture hood and freezing media was aspirated. Then, the brain tissue was transferred to a 2-ml ultra-low attachment vial. The vial was filled with 1 ml of cold hESFM media and the tissue was homogenized with tweezers for 1 minute. This step was followed by subsequent manual homogenization with hand pipettes, first with albumin-coated 1-ml pipette tips and then 200-μl albumin-coated tips. Finally, the homogenized brain tissue was microcentrifuged at 200 rcf for 5 minutes, and the supernatant was aspirated and discarded. The vial was then filled with cold hESFM media to transfer the tissue to the filtration device.

### Filtration Device Design and Microvessel Enrichment

A custom 3D printed filtration device was developed to enrich microvessels from homogenized tissue. Designs for the filtration device are available on Github. As previously described, the 3D printed parts are coated with parylene before use to ensure biocompatibility. The device contains three components: a upper cap with threads and a lip, a lower cap, and a laser cut nylon mesh (100 μm filter size) in between. The nylon mesh fits in the center of the bottom cap, and then the top cap is screwed onto the lower cap. A gasket o-ring (McMaster Carr 90025k364) is placed underneath the lip on the top cap to act as a seal. The entire filtration device is then placed in a 150-ml side arm erlenmeyer flask (Fisher 10-180D). The side arm is then attached to a pipette aspirator. Using a pipette tip, the homogenized brain tissue is transferred from an ultra-low attachment vial onto the mesh filter using an albumin-coated pipette tip. Sterile PBS is then poured onto the mesh while slightly pulling pressure with the pipette aspirator. This strategy pulls the PBS down through the filter to carry away dead or singularized cells, leaving behind enriched microvessels. After this step, the mesh filter is removed and the microvessels are gently scraped into an ultra-low attachment 6-well plate containing cold hESFM for additional washing. Next, the microvessels are transferred to a 2-ml ultra-low attachment vial and centrifuged at 400 rcf for 5 minutes. Last, the supernatant is aspirated and the microvessels are resuspended in 250 μL of 10% GelCad precursor solution (w/v in complete media) containing 15 μL microbial transglutaminase (Modernist Pantry Moo Gloo TI 1203-50).

### Microvessel culture in microfluidic chips

After microvessels have been isolated and resuspended in GelCad precursor solution with transglutaminase, microdevices are prepared for injection. First, microfluidic chips are placed in the 3D printed holder with the long part of the central channel face down (Supplemental Figure 1). Using the PDMS fragments originally removed via biopsy punch, the flanking perfusion channels are temporarily plugged at the inlet and outlet, which helps prevent hydrogel from vacating the central channel after injection. An electronic autopipette is then used to inject the microvessel solution into the central channel. The autopipette facilitates a uniform, slow injection that is continued until the microvessel solution reaches the port on the opposite side of the device. After injection, microdevices are carefully removed from the chip holder and placed into an incubator to crosslink for a minimum of 4 hours and a maximum of 18 hours. After hydrogel crosslinking was confirmed, the microfluidic chip was connected to the pump perfusion system. Perfusion experiments were run for 14 days, unless otherwise specified. A complete build of materials is provided on GitHub.

### Immunohistology

For fixation, the hydrogels from the microfluidic chips were cut out of the PDMS as previously described.^17^ The hydrogels were then submerged in 4% paraformaldehyde for 30 seconds. The fixed hydrogels were then washed with PBS 3 times at 15 minute intervals. Next, the fixed hydrogels were permeabilized in PBS containing 0.1% Triton X-100. 0.1% Sodium Azide and 5% normal goat serum overnight at 4°C on a rocker. Hydrogels were subsequently washed 5 times with PBS and then blocked again with PBS containing 0.05% Tween-20 and 5% normal donkey serum for 40-60 minutes. Hydrogels were then incubated with primary conjugated antibodies overnight at 4°C on a rocker. All antibodies were used at a concentration of 1:1000. The antibodies used were as followed: NG2-Alexa Fluor 647 (Abcam; ab183929), Aquaporin 4-Alexa Fluor 488 (Abcam; ab284135), PODXL-Alexa Fluor 568 (Abcam; ab211225), Beta-III Tubulin-Alexa Fluor 555 (Abcam; ab202519), Ki67-Alexa Fluor 488 (Abcam; ab197234), PECAM1 – Alexa Fluor 488 (Cell Signaling; 42777), Alpha-Smooth Muscle Actin eFluor 660 (Thermofisher; 50-9760-82), Claudin 5-Alexa Fluor 488 (Thermofisher; 352588), PDGFRB-Alexa Fluor 488 (Thermofisher; 53-1402-82), Lectin DyLight 488 (L32470), Actin (Phalloidin)-Alexa Fluor 555 (Thermofisher A34055), Collagen IV-Alexa Fluor 647 (Thermofisher; 51-9871-82). Fluorescent images of immunostained structures were obtained using a Zeiss LSM 880 confocal microscope at ×10 and ×20 magnifications. The hydrogel was scanned at various focal planes, spanning from z = 0 to 100 μm, with intervals of 8-10 μm. For the purpose of generating a 3D representation of the images, the 3D Viewer plugin within ImageJ software was employed.

### Measurement of permeability in capillaries

The permeability of a 10 kDa dextran labeled with Texas Red was determined by analyzing the diffusion from the vasculature into the surrounding hydrogel matrix. For each permeability assay, a separate microfluidic chip was used, constituting a biological replicate. To locate capillaries for permeability measurements, we first identified vessels with a diameter range of 5–10 μm. A secondary criteria was that each selected capillary needed to be at least 50 μm away from any other vascular structure to minimize external influences on the diffusion measurement. Fluorescent microscopy was used to capture the initial and final distribution of the dextran within a specified field of view (FOV). The initial FOV was defined as 244.03 μm by 193.99 μm, with image resolution of 1385 by 1101 pixels. Fluorescence intensity was quantified from a 10×10 pixel box within this FOV, averaged, and recorded as a single value corresponding to discrete time points over a duration of 3 hours. Red pixel quantification was carried out on the images, with ‘red’ defined by a custom threshold that discriminates the dextran fluorescence from the background. A pixel was classified as ‘red’ if its red channel intensity exceeded a predefined minimum value and was higher than that of the green and blue channels. The total count of ‘red’ pixels represented the dextran-laden area at each time point.

The permeability coefficient was calculated as follows:

1. Area per pixel: The area represented by each pixel (A_pixel_) was derived by dividing the FOV area by the total pixel count in the image.

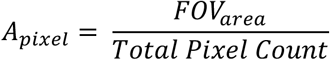
2. Dextran concentration: The initial concentration of dextran (C_0_) was set at 10 mM. This concentration was used to determine the molar quantity of dextran within the red pixel area at the initial and final time points.

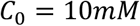
3. Molar flux (J): The flux was calculated using the change in molar quantity of dextran (Δ*n*) over the diffusion area (*A*_*diffusion*_), normalized by the experiment duration (t).

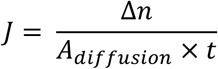
4. Permeability coefficient (P): Fick’s first law was applied to define P, with J divided by the concentration gradient across the capillary (Δ*C*), which was approximated by the initial dextran concentration (C_0_).

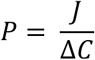

The change in the red pixel count between the initial and final images was used to calculate Δ*n*, while *A*_*diffusion*_ was assumed to be the total red pixel area at the initial time point. P was initially obtained in μm/min and then converted to cm/s for standardization:

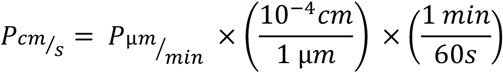

## Results

### Vessel purification and assessments of structural integrity

Extracting intact microvasculature from human tissues is a challenging endeavor due to small size, relative scarcity within tissue, intricate architecture, and the need to preserve cellular viability. Traditional methods often rely on enzymatic digestion, which can lead to partial degradation or loss of critical components such as endothelial cell tight junctions and the arrangement of other cells in and around the vessels, thus compromising the structural integrity of the vessels. To address this challenge, we developed an enzyme-free protocol for the extraction of human microvessels from cryopreserved postmortem cortical tissue, aimed at preserving the structural integrity of the microvascular networks (Figure 2A). After isolation, the enriched microvessels were embedded in a hydrogel within a perfused microfluidic device for 4 days before initial analysis—the hydrogel and microfluidic device are described in more depth below. After 4 days of culture, the microvessels had high viability as seen by calcein staining (Figure 2B). To validate the structural integrity and cellular composition of the enriched vasculature, we performed extensive immunostaining, with the results presented in Supplemental Figure 2. The staining results reveal a comprehensive picture of the preserved cellular architecture. We provide evidence of both capillary and arteriole structures based on vessel size and the presence of NG2+ and α-SMA+ mural cells around PECAM-1+ and claudin-5+ endothelial cells. Positive aquaporin-4 (AQP4) signal around vessel structures suggests the presence of astrocyte endfeet, and immunostaining for PODXL, a sialoglycoprotein found on the luminal side of brain endothelial cells,^18^ indicates maintenance of the glycocalyx. We could detect βIII-tubulin+ neurons that are likely remnants of the purification process, but we did not assay for their survival or include them in any downstream analyses. Last, positive immunostaining for Ki-67 suggests vessel growth was occurring after embedding in the hydrogel. Overall, these data support our claims that the enzyme-free extraction method effectively preserves the structural integrity of isolated human microvessels and enables their culture within hydrogels.

**Figure 2:**
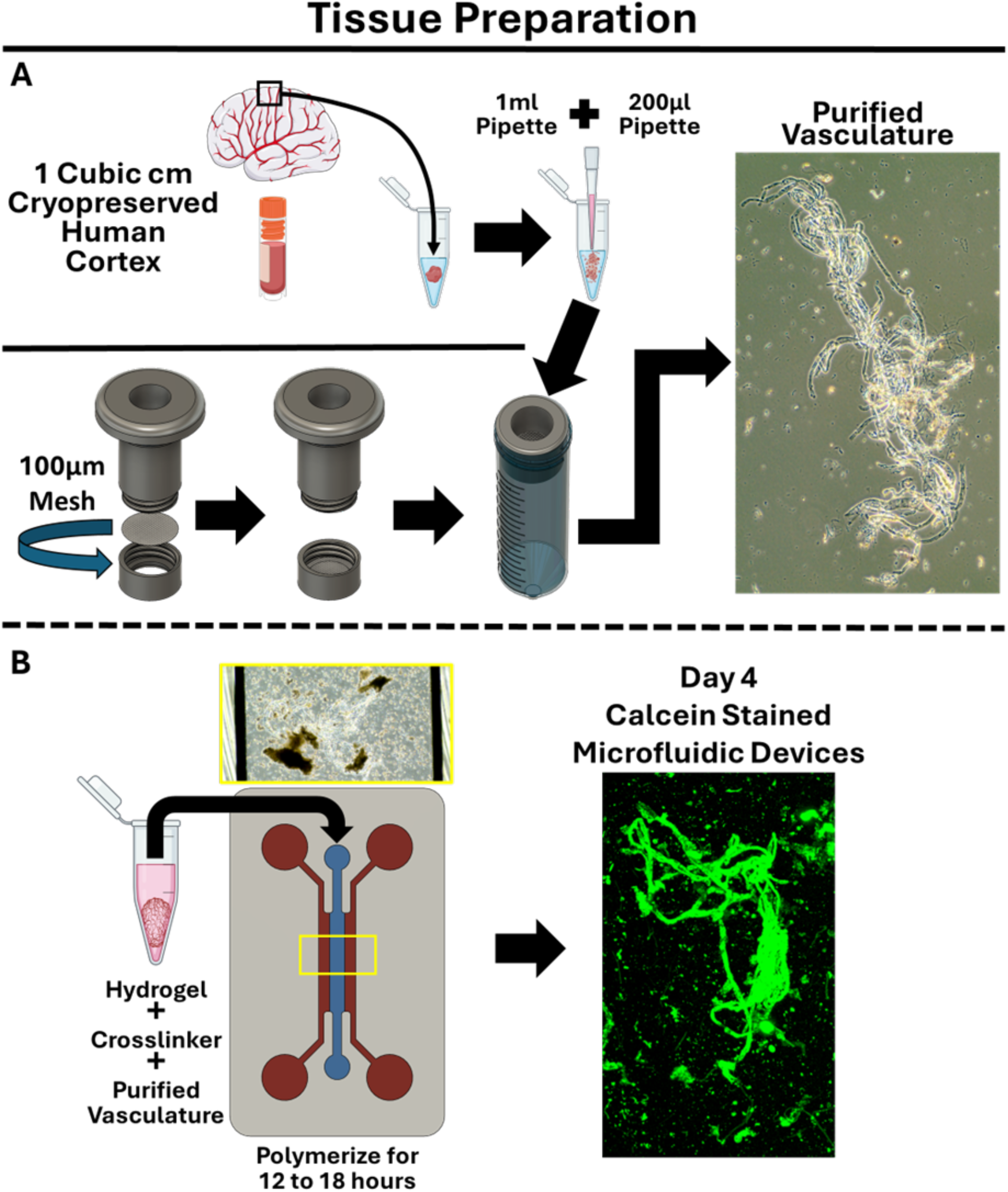
Strategy for obtaining and culturing purified vasculature from cryopreserved human cortical tissue. **A)** ∼1 cm^3^ piece of cryopreserved human cortical brain tissue is transferred to a 2-ml ultra low attachment conical tube and homogenized using hand pipettes. Vessels are purified from the dissociated tissue using a custom filtration device equipped with a 100 μm mesh filter. A representative brightfield image of isolated vessels is shown on the right. **B)** Purified vessels are reconstituted in a hydrogel precursor solution mixed with an enzymatic crosslinker. The mixture is gently injected into the center chamber of the custom microfluidic device, allowing the hydrogel to crosslink overnight. A representative image of calcein-stained tissue is shown on day 4 of culture within the microdevice, showcasing high viability of the microvessels.

### Pump perfusion system for recirculating media and pulsatile wave generation

After confirming structural integrity of isolated human microvessels with hydrogels, we aimed to establish extended culture of the human microvessels, accounting for several factors. One essential factor was the ability to deliver fresh media to the embedded vasculature for nutrient and waste product exchange. Typically, in a microfluidic system setup, a reservoir of media is attached to a pressure pump to consistently deliver fresh media to the embedded cell or tissues in the microfluidic devices. These devices use a low volume of media and are easily controlled using commercially available systems. However, because the media is not recirculated, cells are unable to create the optimal conditions in their microenvironment. Cells condition their surrounding media over time, which alters the balance of nutrients and metabolic byproducts in the media. Moreover, cells secrete signaling molecules such as extracellular vesicles, cytokines, growth factors, and ECM proteins, which can influence both their own behavior and that of neighboring cells via paracrine and autocrine signaling.^19^ Due to these factors, we believed that a recirculating fluid system would not only maintain continuous exchange of culture media, but also enable extended culture by addressing critical aspects of media conditioning. Additionally, due to the nature of the isolated vessels containing arterioles, we believe that a hemodynamic force would be necessary for arteriole growth. Extensive research has established that the circumferential mechanical strain of arteries and arterioles is a crucial element in vascular growth, and plays a critical role in regulating vascular structures and function.^20^ However, to our knowledge there are no commercially available pump perfusion systems that are able to mimic the endogenous arterial pressure wave, including the characteristic dicrotic notch. To address this gap, we developed a custom pump perfusion system capable of generating physiologically accurate pulse waves to the microfluidic systems and allow meticulous control over pulse rate, pressure wave amplitude, pulse frequency, dichroitic notch timing, and pulse width modulation (Figure 3). Using open-source hardware, and a free custom graphical user interface simplifies the creation of pulse waves tailored to specific experimental needs, further enhancing the usability of this system for our desired application and beyond.

**Figure 3:**
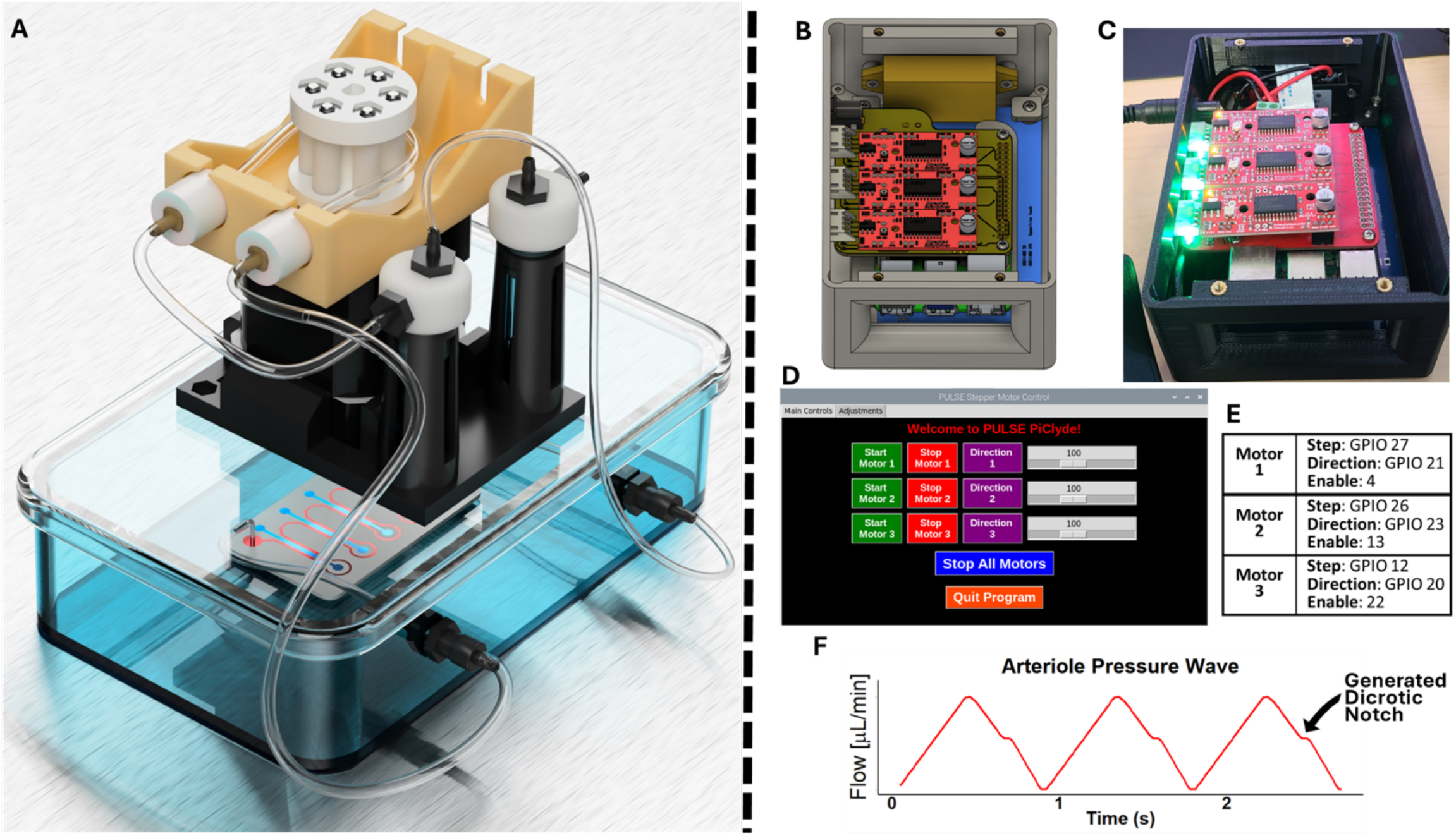
Custom pump perfusion system. **A)** Representation of the experimental system, showing the stepper motor constituting the pump, the container holding the microfluidic device, the device itself, and the fluid reservoir. **B)** CAD design of the custom pump perfusion system, illustrating the EasyDriver motor controllers, custom PCB, Raspberry Pi, touchscreen, and housing unit. **C)** Actual built design of the CAD design, showcasing the assembled components. **D)** Custom graphical user interface developed to control three motors individually, providing precise control over each motor’s operation. **E)** Stepper motor with corresponding step, direction, and enable pins, essential for controlling the motor’s movements. **F)** Pulse wave generated by the pump perfusion system, as recorded using a Sensirion flow sensor. The pulse wave includes a dicrotic notch, indicated on the graph, which is present in arterial pulse waves.

### Vascularization of hydrogel-laden microfluidic devices under constant perfusion

Next, we proceeded with extended culture of the human microvessels within hydrogel-laden microfluidic devices. We used a gelatin-based hydrogel functionalized with a biomimetic peptide derived from the homophilic cell adhesion epitope of N-cadherin (“GelCad”) to promote cell integration in the 3D environment. Previous work has shown that the conjugation of N-cadherin peptides with biomimetic hydrogels are largely beneficial for cellular integration in 3D environments.^15,16,21–24^ Hence, we anticipated that GelCad hydrogels would support survival and outgrowth of human microvessels. To this end, we cultured microvessels for 14 days in the microfluidic devices under constant recirculating perfusion. Immunostaining at the endpoint revealed robust networks of claudin-5+/collagen IV+ vessels (Figure 4A). Visualization of lectin and αSMA further revealed the presence of αSMA+ arterioles connected to αSMA-/lectin+ capillary-like structures (Figure 4A). Imaging of the lectin+ vessels near the edge of the hydrogel demonstrated anastomosis with the perfusion channel (Figure 4B), indicative of fluid entry into the vessels from the perfusion channel, which we explored later.

**Figure 4:**
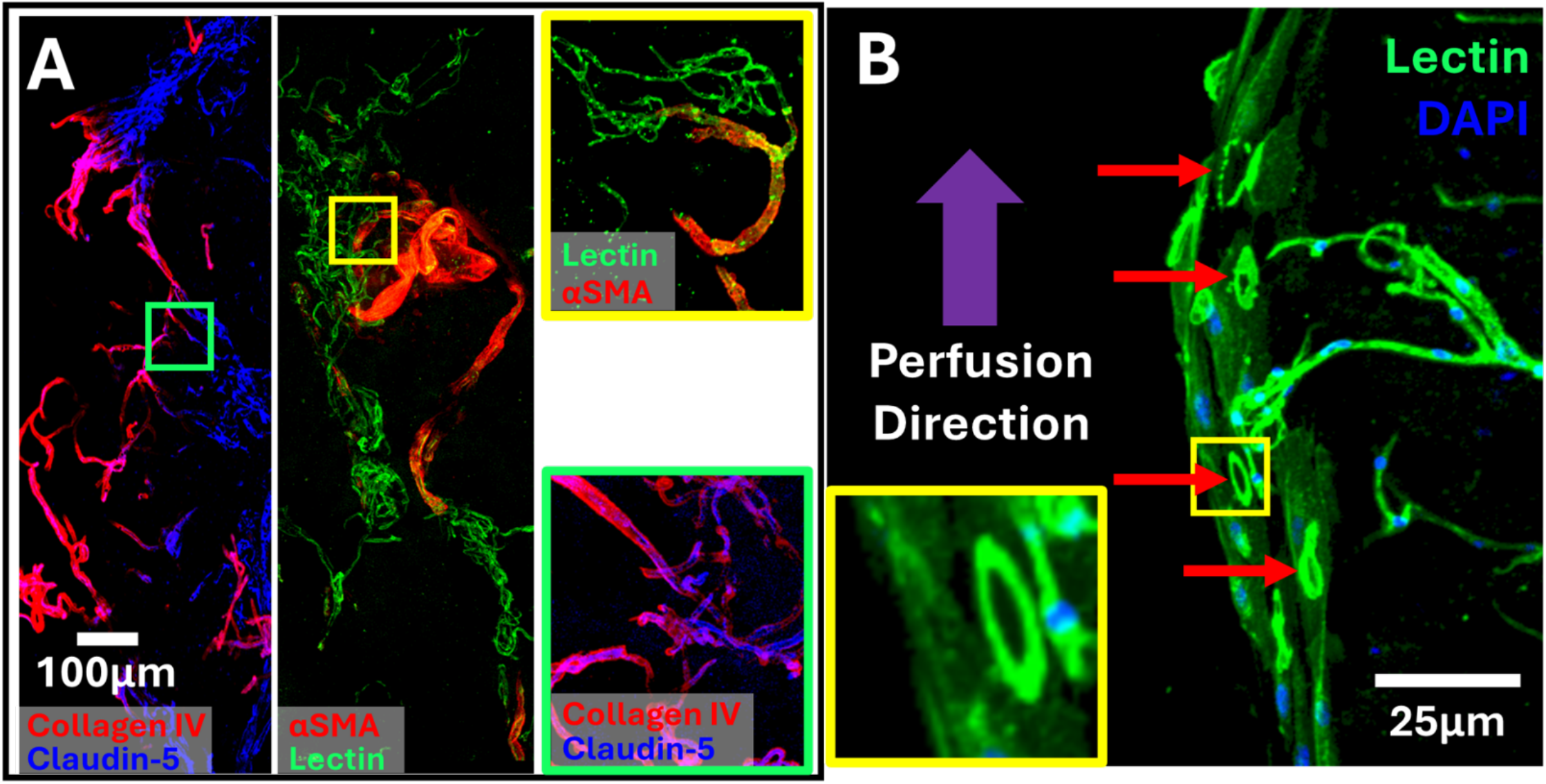
Vascularization of hydrogels within the microfluidic device after 14 days of culture. **A)** Representative images of collagen IV and claudin-5 (left panel), and αSMA and lectin (right panel). The yellow inset highlights the branching of αSMA+ arterioles down into lectin+ capillaries. The green inset features extensive vascularization within the hydrogel. **B)** Representative images of lectin+ vessels at the interface of the hydrogel and the perfusion channel. The red arrows and yellow inset highlight the formation of lumenized structures along the edge of the hydrogel.

More extensive immunostaining at the 14-day endpoint confirmed robust vessel architectures. We were able to detect arterioles with *in vivo*-like architectures consisting of concentric rings of smooth muscle cells wrapping around claudin-5+ endothelial cells, separated by a collagen IV+ basement membrane (Figure 5A). Cross-sectional images showcased the patent lumens of these arteriole structures (Figure 5A, panels i and ii). We could further detect conjoining claudin-5 tight junctions within the collagen IV+ structures and NG2+ pericytes wrapping around the outside of capillary-sized vessels (Figure 5B-C). AQP4 signal remained detectable on the outside of vessels, again suggesting the presence of astrocyte endfeet (Figure 5D). Collectively, these results demonstrate that the engineered human microvessels can be effectively grown within hydrogel-laden microfluidic devices, accurately replicating the spatial arrangement and organization of the NVU and BBB, mirroring the native vessel architecture of the human brain.

**Figure 5:**
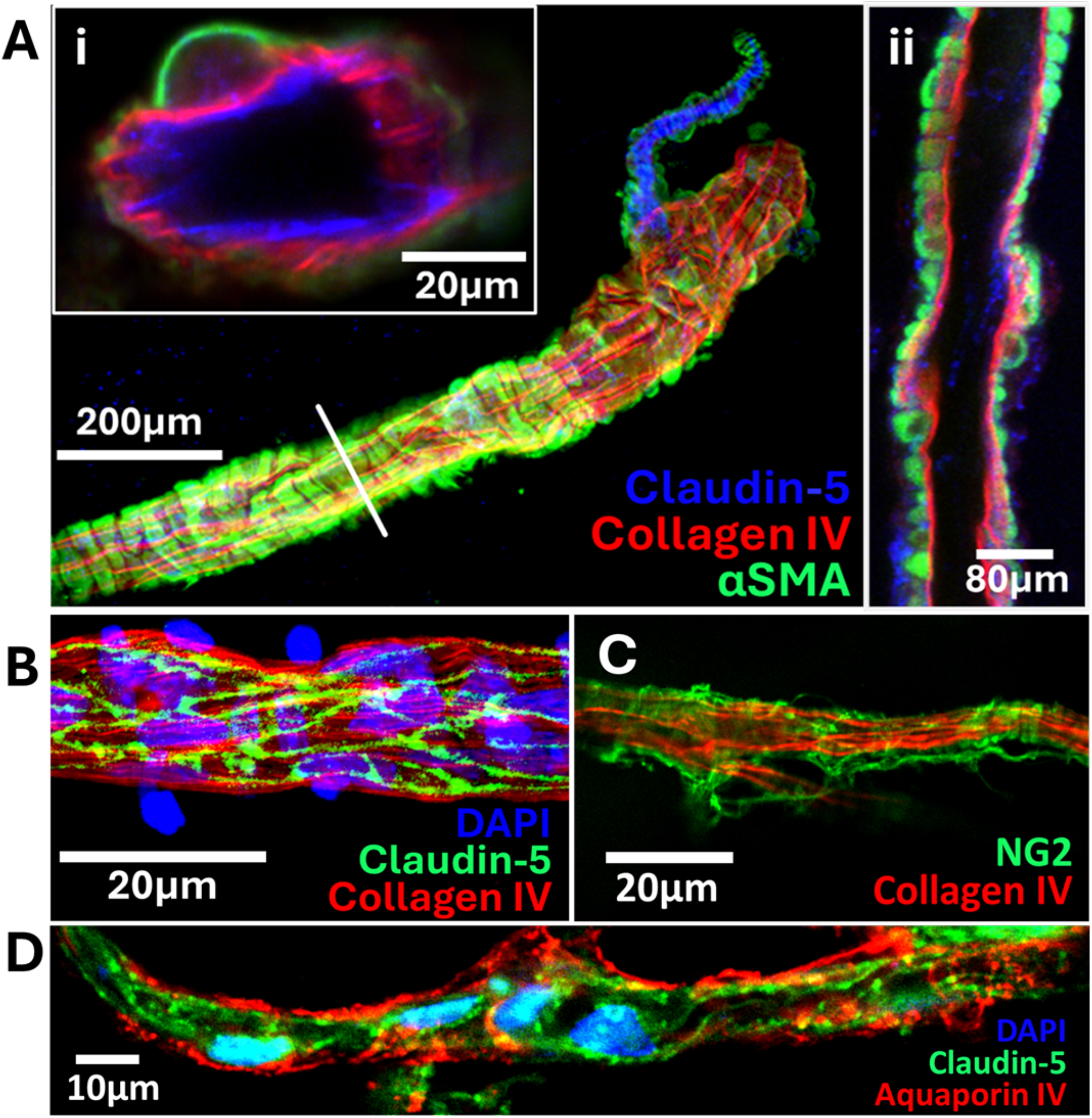
Anatomical arrangement of arterioles and capillaries within the microfluidic device after 14 days of culture. **A)** Representative image of an arteriole identified by concentric rings of smooth muscle cells. Insets show different cross-sectional images highlighting the correct concentric anatomical arrangement of the arterioles. **B-C)** Representative images of capillaries showing intact tight junctions in endothelial cells (panel B) and pericytes wrapping around the vessel (panel C). **D)** Representative image of AQP4+ endfeet surrounding a vessel. Each panel of this study is from an individual biological replicate (n=4).

### Evaluation of engineered microvessel perfusion and BBB permeability

One of the most critical challenges in developing *in vitro* models of the BBB is accurately assessing the permeability and integrity of the engineered barrier. The BBB’s ability to selectively permit the passage of molecules while restricting harmful substances is essential for maintaining CNS homeostasis. To evaluate the permeability of microvessels in our system, we utilized a 10 kDa dextran conjugated to Texas Red dye. At our standard 14-day timepoint, we introduced the dextran into the flanking perfusion chambers, where it readily entered anastomosed vessel lumens indicating their ability to be perfused (Figure 6A). Time-lapse confocal microscopy was then used to assess dextran extravasation from the lumen of individual capillaries to the extracellular matrix. The baseline image taken at 0 minutes (Figure 6B) shows the initial distribution of dextran. After 180 minutes (Figure 6C), the dextran remained largely confined within the vascular lumen, with only minor diffusion into the surrounding tissue. At the end of the assay, immunostaining was used to confirm capillary integrity by the presence of claudin-5+ endothelial cells and a collagen IV+ basement membrane lacking coverage by smooth muscle cells (Figure 6D-E). The quantitative analysis of BBB permeability, presented in Figure 6F, showed a permeability coefficient for Dextran was calculated to be 1.154e-07 cm/s, a value consistent with those observed in other BBB models.^8,12,17^ These results are particularly significant when compared to other *in vitro* models, which often struggle to maintain such low levels of permeability, possibly due to the lack of dynamic flow conditions and proper anatomical arrangement.

**Figure 6:**
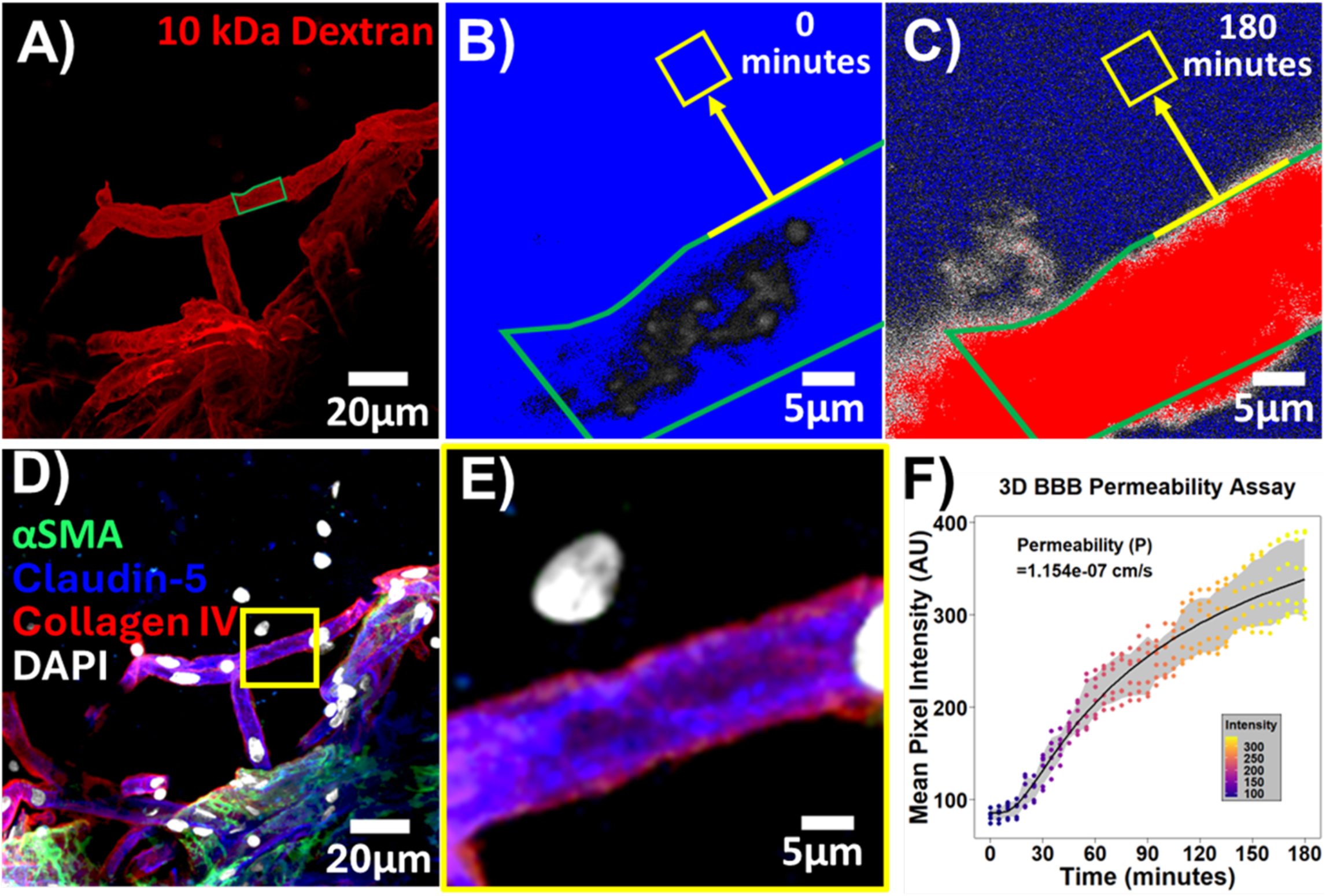
BBB permeability assay. **A)** Representative image of Texas Red-labeled 10 kDa dextran perfusion through vessels at the day 14 time point. **B-C)** Example baseline image (time = 0 minutes) and final image (time = 180 minutes) for dextran extravasation. The yellow box indicates the specific region of interest, taken 10 μm away from the capillary, for the full time-lapse imaging. **D-E)** Immunofluorescence labeling of the fixed vessels after the permeability assay to identify and characterize the cellular components of the vasculature. The yellow inset highlights the region where the time-lapse imaging was performed. Comprehensive labeling confirms the assay was performed in a capillary lacking αSMA coverage. Quantitative analysis of BBB permeability based on the change in Texas Red signal over time. Measurements were collected every 5 minutes, and each dot at the respective time point represents an individual biological replicate (N=6 independent microdevices). The calculated permeability coefficient (P) is 1.154e-07 cm/s, and the shaded region represents the standard error of the mean (SEM) from the biological replicates.

## Discussion

In this study, we demonstrate the ability to engineer lumen-perfusable, anatomically accurate, and physiologically relevant human cerebrovasculature within a microfluidic device. The intricate composition of the cerebrovasculature, involving tightly coordinated interactions between endothelial cells, mural cells, astrocytes, and the ECM, is essential for maintaining CNS homeostasis and BBB integrity. However, to our knowledge, there are currently no methods that have effectively grown *ex vivo* vasculature with the correct architecture to represent both the NVU and BBB. Our new technology addresses several longstanding obstacles in replicating these complex structures *in vitro*. To overcome challenges associated with promoting self-assembly of individual neurovascular cell types into vessel-like structures, we employed the novel approach of culturing purified human cortical microvessels in a biomimetic hydrogel connected to a custom perfusion system. This system promoted growth and anastomosis of vessels to the edges of the hydrogels, and we could detect perfusable vessels with both arteriole and capillary architectures that were interconnected in the hydrogel.

Most models of the cerebrovasculature have relied on microfluidic device designs that enable culture of brain endothelial cells on porous planar substrates or on the edges of hydrogels, which can be further interfaced with a separate chamber harboring neural or glial cells.^10,25–30^ In contrast, engineered models of three-dimensional vascular networks at the level of capillaries and arterioles generally rely on self-assembly of individually prepared cell types into representative vascular structures within hydrogels. Some notable model examples include the assembly of endothelial cells, pericytes, and astrocytes into perfusable vessel structures in microfluidic devices,^8,9,17^ and the assembly of endothelial cells, mural cells, and astrocytes under static conditions.^31^ However, these self-assembled models have noted structural deficiencies that our use of primary human microvessels can overcome. As one example, to our knowledge, no prior models constructed with individual cells have demonstrated concentric rings of smooth muscle wrapping around lumenized endothelial cells, representing bona fide arteriole structures. Generally, it has also been challenging to achieve vessel diameters representing capillary formation (5-10 μm), and recent advancements that can achieve endothelialized structures of this size still lack pericyte coverage.^32^ Astrocytes, while present in some of these systems, do not typically extend their endfeet along the entirety of the vessel surface,^12^ as seen *in vivo*. Further, arteriole and capillary vessels have not been generated within the same hydrogel, nor with interconnected perfusion in the same vascular network. We provide evidence that each of these features can be achieved with primary human microvessels cultured in our biomimetic hydrogel. We believe that a critical aspect of our approach is the ability to preserve microvasculature integrity and cell viability through an enzyme-free extraction method. This step maintains cerebrovascular architecture within the microfluidic platform as a starting template for vascular proliferation. The preservation of these structures is vital for studying the NVU, where cell-cell and cell-matrix interactions are more likely to mimic native *in vivo* conditions. As such, our model presents a valuable tool for advancing research into disease mechanisms and therapeutic testing in a physiologically relevant environment.

After replicating the structural characteristics of the NVU, we assessed permeability of capillaries using a dye extravasation assay. Unlike many models that fail to achieve the physiological tightness of the BBB, our *ex vivo* model maintains a low permeability coefficient comparable to *in vivo* conditions.^33^ Comparisons between planar versus more lumenized *in vitro* BBB models, particularly those fabricated from iPSCs, suggest three-dimensional architecture is important for maintaining robust BBB function.^11,26,29^ In our system, the close apposition of pericytes to the brain endothelial cells likely reinforces BBB function.^34,35^ Beyond cellular organization, a unique feature of our model is the application of a hemodynamic profile within the microdevices that mimics the arterial pulse wave, including the dicrotic notch. Hemodynamic forces are known to play an important role in vascular growth, remodeling, and integrity.^36,37^ Future work will focus on assessing how such hemodynamics influence salient features of our model system.

In conclusion, our study presents a significant advancement in the field of neurovascular research by successfully engineering physiologically relevant NVU and BBB organization and function within a microfluidic platform. The accurate replication of the NVU’s complex spatial organization and the physiological properties of the BBB could provide a valuable tool for studying neurovascular diseases and drug delivery across the BBB. The combination of human cerebral blood vessels, biomimetic hydrogels, and innovative microfluidic and perfusion technologies allows for the creation of a model that closely mimics the *in vivo* environment, offering a bridge between *in vitro* studies and clinical applications. One limitation of our model is the requirement for postmortem donor tissue, but our use of cryopreservation enables model scalability limited only by the number of banked tissue samples. This proof-of-concept study utilized a single donor for all experiments, and future work will focus on expanding these analyses across multiple sources of donor tissue.

## Acknowledgments

Funding for this work was provided by Ben Barres Early Career Acceleration Award from the Chan Zuckerberg Initiative (grant 2018-191850 to ESL) and NIH grants R01 NS110665 (to ESL), RF1 NS130334 (to ESL and MSS), and K99 NS133399 (to BJO). BJO was also supported by the Vanderbilt Interdisciplinary Training Program in Alzheimer’s Disease (T32 AG058524). ASM was supported by the Interdisciplinary Training Program in Lung Research (T32 HL094296). AK was supported by an NSF Graduate Research Fellowship. Some image acquisition and analyses were performed through the use the Vanderbilt Cell Imaging Shared Resource (supported in part by P30 CA068485, P30 DK058404, and P30 EY008126).

## Availability statement

All designs and files from this work are available at: https://github.com/OGradyLab/NVU-On-A-Chip.

## Figures

**Supplemental Figure 1:**
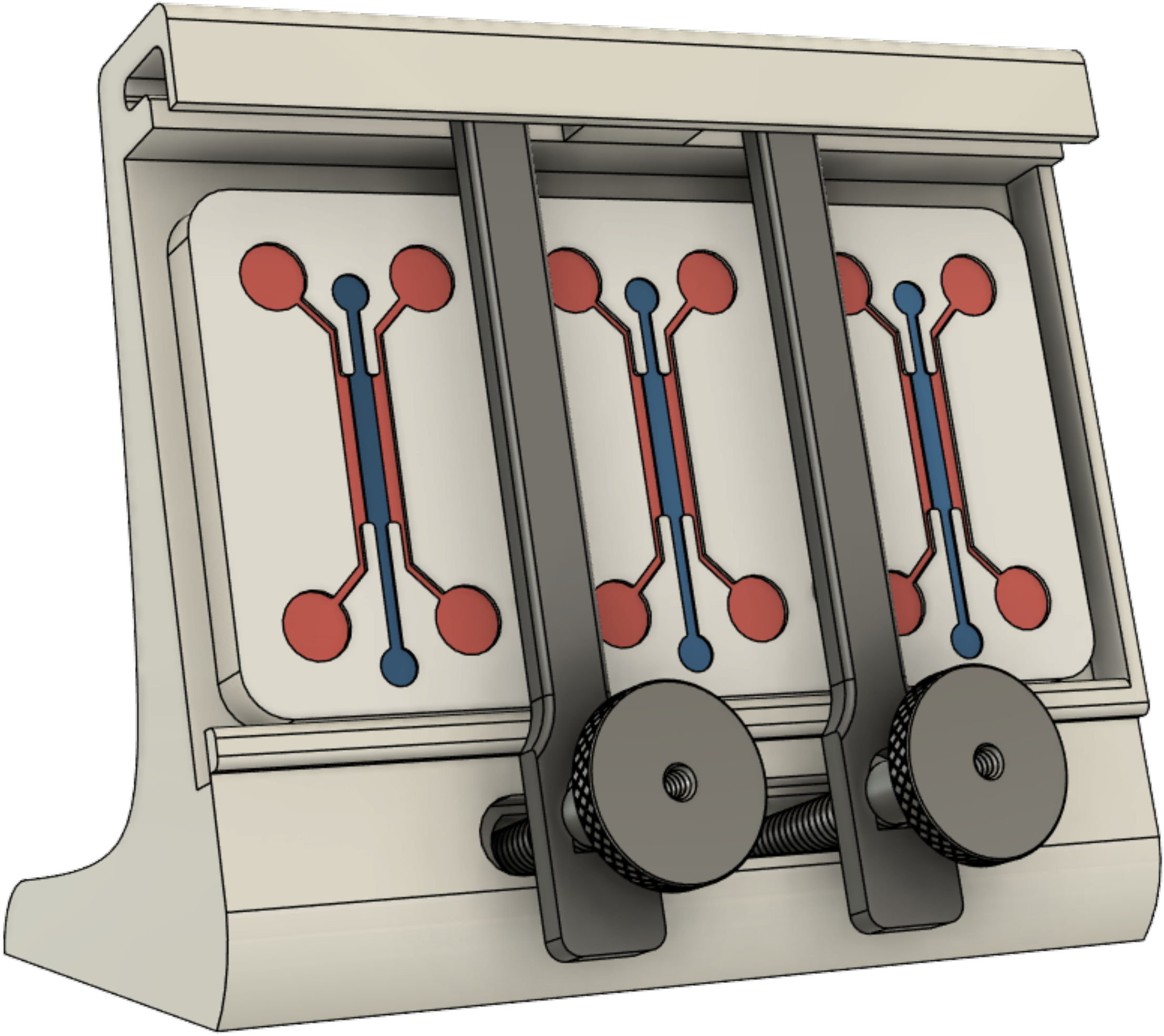
Custom microfluidic chip holder designed for hydrogel injection. The holder positions the chip at a precise angle to ensure uniform filling of the central channel with hydrogel while preventing spillage into adjacent perfusion channels.

**Supplemental Figure 2:**
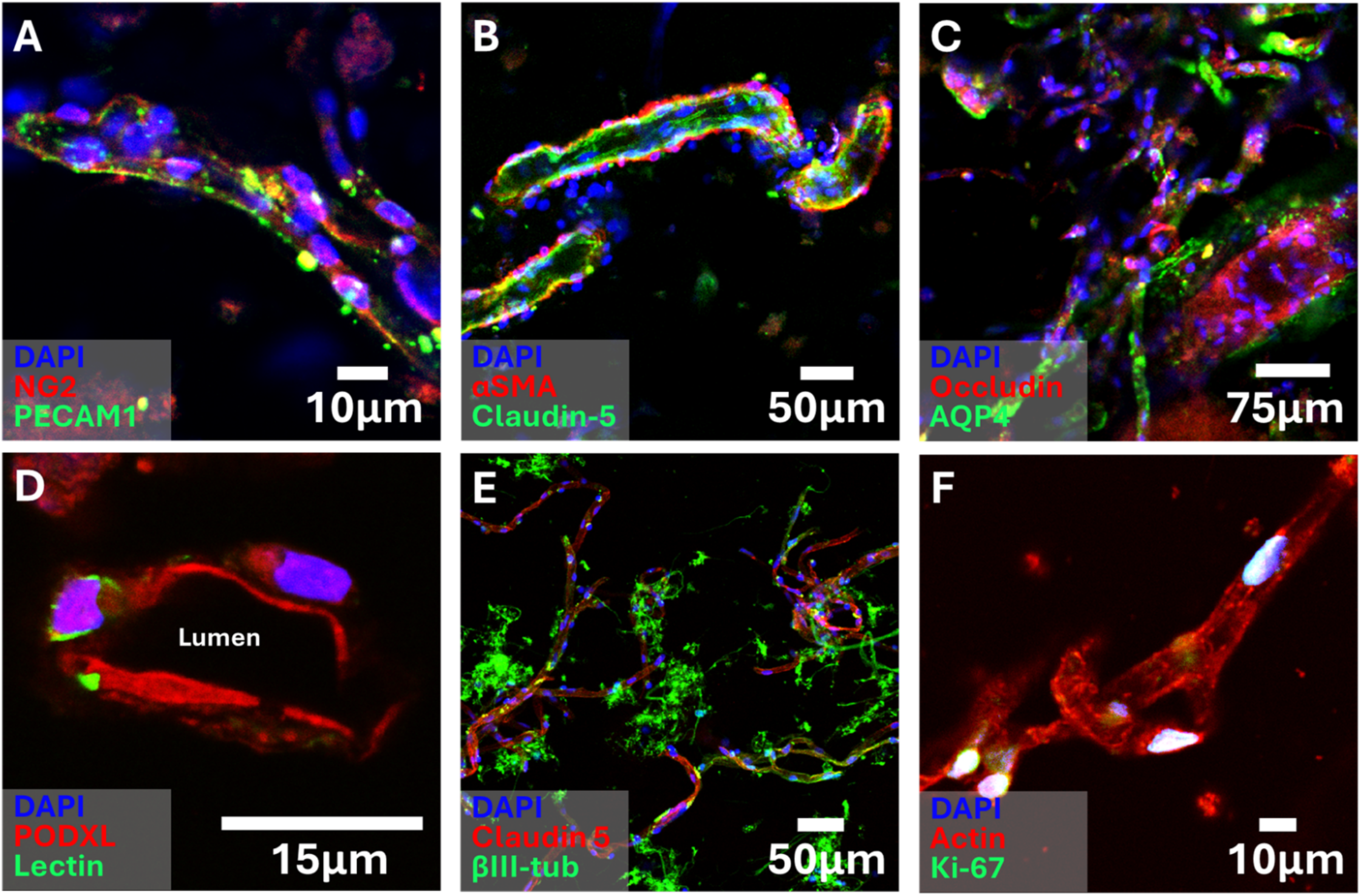
Immunofluorescence staining of microvessels cultured within the hydrogel for 4 days. **A)** Endothelial cells (PECAM1, green) and pericytes (NG2, red) forming intact microvessel structures. **B)** Smooth muscle cells (αSMA, red) surround the endothelium, confirming the presence of arteriole-like structures, while tight junctions (Claudin-5, green) are clearly visible. **C)** Aquaporin-4 (AQP4, green) highlights astrocyte endfeet coverage around vessels. **D)** Lectin (green) binds to endothelial glycocalyx, confirming lumenized structures, with PODXL (red) marking the endothelial surface. **E)** βIII-tubulin+ neurons (red) are visible near Claudin-5+ (green) microvessels as potential remnants of the vessel isolation procedure. **F)** Proliferating cells are marked by Ki-67 (green), with actin (red) revealing vessel structure.

